# Predator type shapes call-type composition and post-threat vocal dynamics in Japanese quail (Coturnix japonica): a pilot case-series

**DOI:** 10.64898/2026.09.11.750852

**Authors:** Kayeon Ham, Seeun Park, Goun Park, Chunghyun Kim

## Abstract

Japanese quail (Coturnix japonica) rely on an extensive vocal repertoire to coordinate social behaviour and respond to predation risk, yet how individual call types are modulated by predator identity remains poorly characterized. In this pilot case-series, two sexually mature Japanese quail (each contributing more than 2,000 classified vocalizations across repeated predator-exposure trials) were exposed to simulated aerial (hawk) and terrestrial (cat) predator stimuli across four social contexts, before, during, and after stimulus presentation. Vocalizations were classified, via blind dual-observer review, into 18 spectrographically distinct call types (an exploratory repertoire estimate, not independently statistically validated), of which 10 were acoustically characterized. Chi-square analyses of the full call-type composition revealed highly significant differences between hawk and cat exposure in both individuals (both p < 0.0001); call types 5 and 10 were significantly cat-biased and call type 2 was significantly hawk-biased in both individuals. An exploratory correlation suggested that call types with a higher maximum frequency tended to be relatively hawk-biased (Spearman rho = -0.67, p = 0.033, uncorrected for multiple comparisons). Independently, call-type composition shifted significantly between the pre- and post-exposure phases in all four individual-by-predator combinations (all p < 0.0001), driven substantially by a consistent post-exposure increase in call type 9 across both individuals and both predator classes. Additional exploratory analyses of social-context (mirror and concealed-conspecific) effects on call-type composition are presented in the supporting information. Given the small sample size (n = 2), these findings are presented as case-based, hypothesis-generating observations intended to justify and inform a fully powered follow-up study, rather than as population-level conclusions.

## Introduction

Domestic and wild Galliformes possess vocal repertoires that encode information about predator class, food availability, and social context [1,2]. In the domestic chicken (Gallus gallus domesticus), playback experiments have established that acoustically similar call types can nonetheless carry distinct, functionally referential information: food calls and ground-alarm calls share comparable pulsatile acoustic structure yet elicit categorically different receiver responses [3], and alarm calling itself is modulated by the presence and identity of an audience [4,5].

Vocal repertoire size and structure have been quantified across a range of avian taxa, including Korean passerines such as the Daurian Redstart (Phoenicurus auroreus), for which spectrographic classification has revealed an extensive song repertoire structured into distinguishable component parts [6]. The Japanese quail (Coturnix japonica) is reported to possess a comparably large vocal repertoire, historically described as comprising approximately two dozen discrete call types organized into four functional groups, including one linked to strong defensive responses toward interspecific threats such as predators [7]. Individual call types, most notably the male crow, have been examined for their role in mate recognition and subspecies discrimination [8,9], and isolation-induced distress calling has been shown to be suppressed by visual social cues, including mirror images [10]. In the domestic chicken, an ecologically embedded mirror-audience paradigm, in which alarm calling toward a simulated aerial predator was compared across solitary, conspecific, and mirror conditions, found that roosters warned a live conspecific but not their own mirror reflection, even when a real conspecific was concealed behind the mirror [11]; notably, the same birds failed a classic mark test, illustrating that mirror-related behaviour can differ substantially depending on whether it is assessed ecologically or via the classic self-recognition paradigm. However, we are not aware of a study that has systematically classified the full vocal repertoire of Japanese quail against predator class (aerial versus terrestrial) or examined how call-type composition changes across the pre- and post-threat period; even the interspecific defensive-call group described for European quail was not further differentiated by predator identity. This represents, to our knowledge, an unexamined gap for this species.

Our laboratory has previously used a simulated-predator paradigm (moving hawk and cat silhouettes presented in a dedicated arena) to demonstrate that rearing environment shapes separation-calling frequency in Japanese quail under threat [12]. Building on this apparatus and paradigm, the present pilot study asks whether specific, spectrographically defined call types within the broader quail vocal repertoire are differentially recruited depending on predator class, whether call-type composition changes once a threat has passed, and whether any such pattern is reproducible across individuals despite the well-documented individual variability of galliform vocalizations [8].

## Materials and methods

### Ethics statement

All experimental procedures were approved by the Institutional Animal Care and Use Committee (IACUC) of Hoseo University (approval numbers HSUIACUC-24-039, approved 23 December 2024, and HSUIACUC-25-014, approved 19 May 2025), consistent with the approval framework established for simulated-predator exposure research in this species and laboratory [12]. Predator stimuli were presented as pre-recorded silhouette projections and matched audio cues rather than live predators, and no physical contact between test subjects and any predator stimulus occurred at any time. Behaviour was continuously monitored via the wall-mounted video camera described above throughout each session, and no injury, mortality, or other adverse welfare event was observed in either subject during testing. Reporting follows the ARRIVE guidelines for animal research [13] where applicable to the scale of this pilot case-series.

### Subjects

Two sexually mature female Japanese quail (individual identifiers 0.0.0 and 795), 6 weeks of age, were purchased from Join Farms Co., Ltd., a quail farm located in Eumseong-gun, Chungcheongbuk-do, Republic of Korea. Both individuals had prior experience with the testing arena across all experimental sessions but no prior exposure to mirrors before the study began.

In accordance with the STRANGE framework [14], we report the following sample characteristics. Both individuals were female and 6 weeks of age at the time of testing, sourced from a single commercial quail farm (Join Farms Co., Ltd., Eumseong-gun, Chungcheongbuk-do, Republic of Korea). Prior to testing, both individuals were housed under standard commercial stocking conditions typical of Korean quail farms, in cages measuring 50 x 40 cm arranged in a seven-tier battery system, with approximately 30 birds per cage. Social composition prior to testing (i.e., the specific identities of cage-mates), capture method, and experimental history beyond the present apparatus are not yet fully documented in this draft.

### Apparatus

Testing was conducted in an enclosed arena (2.0 x 1.2 x 0.8 m, L x W x H) constructed of waterproof plywood with a green PVC floor and a mesh-net top (8 cm mesh size), located centrally within a dedicated experimental room fitted with blackout curtains and non-flickering illumination to minimize uncontrolled visual and acoustic stimuli, following the design established for simulated-predator exposure in this species and laboratory [12]. Predator stimuli were presented as moving silhouettes projected onto a 1.4 x 1.0 m white panel suspended from the ceiling above the arena, accompanied by matched auditory cues; a hawk silhouette represented an aerial predator and a cat silhouette represented a terrestrial predator. Behaviour was recorded using a wall-mounted video camera positioned to capture the entire arena, and vocalizations were recorded separately via a microphone for subsequent acoustic analysis [12].

### Predator stimulus presentation and social context manipulation

Each individual was tested under two predator conditions (hawk, cat) and four social-context conditions: (A) alone, (B) exposure to a mirror image of the self, (C) exposure to a familiar live conspecific housed visibly on the opposite side of a transparent partition, and (D) combined exposure to a mirror image together with a live conspecific housed behind a visually opaque, sound-permeable partition, such that only auditory cues from the conspecific were available in this condition. Each experimental session consisted of two repetitions of a three-phase sequence: a 5-minute pre-exposure phase, a 5-minute during-exposure phase, and a 5-minute post-exposure phase, yielding approximately 15 minutes of observation for each phase category per session.

Vocalizations were quantified separately for each phase within each predator-by-social-context combination. Extended analyses of conditions B, C, and D are reported in the supporting information (S1 File); the main text focuses on the pooled predator-type effect (conditions A-D combined) and its temporal dynamics.

### Acoustic recording and call classification

Vocalizations were recorded and analyzed spectrographically using Raven Pro (Cornell Lab of Ornithology, Ithaca, NY, USA), following the call-classification approach used by Lee and Sung [6] for the spectrographic analysis of avian vocal repertoires in South Korea. Call types were first identified by visual inspection of their morphological, temporal, and frequency distinctiveness on the spectrogram, and this classification was subsequently confirmed by blind review from a second, independent observer, following the general procedure described in [6]. Eighteen spectrographically distinct call types were identified from the full recording set. For ten of these call types, sufficient exemplars were available to characterize their acoustic structure quantitatively, including call duration, highest frequency, lowest frequency, frequency range, and dominant frequency (mean ± SD).

### Statistical analysis

Given the small number of individuals (n = 2), population-level inferential statistics (e.g., mixed-effects models with individual as a random effect) were not appropriate. Chi-square tests of independence were used to test whether call-type composition (i) differed between hawk and cat exposure and (ii) differed between the pre- and post-exposure phases, separately for each individual; standardized residuals identified which call types drove any overall difference. Correlations between acoustic parameters and predator-type bias were assessed with Pearson and Spearman coefficients across the ten quantified call types. The threshold for statistical significance was set at alpha = 0.05 for all tests unless otherwise specified; the acoustic-correlate analysis (five parameters tested) was additionally evaluated against a Bonferroni-corrected threshold of alpha = 0.01 to account for multiple comparisons, as reported in Results. Cross-individual reproducibility of directional patterns was assessed qualitatively as convergent case-based evidence rather than through population-level null-hypothesis testing.

## Results

### Overall calling frequency and cross-individual convergence by predator type

Total call counts (summed across all social contexts and temporal phases) for each of the ten quantitatively characterized call types are shown for both individuals in Table 1, together with the direction of the cat-versus-hawk difference for each individual and its consistency across individuals.

**Table 1.**
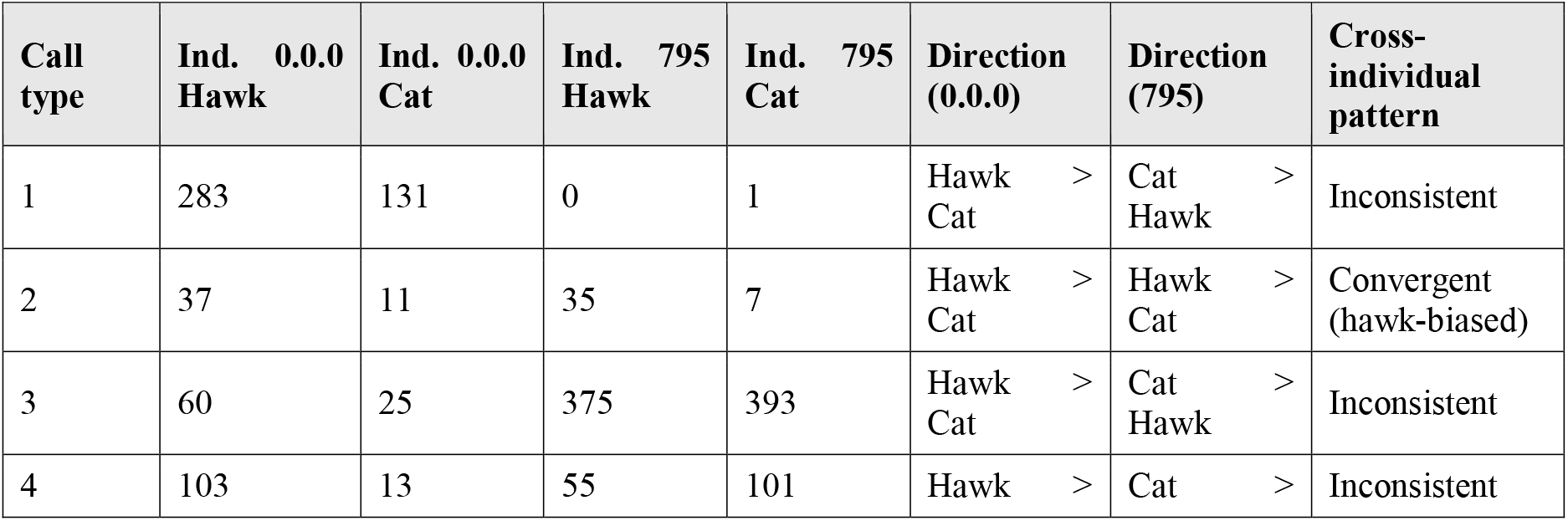
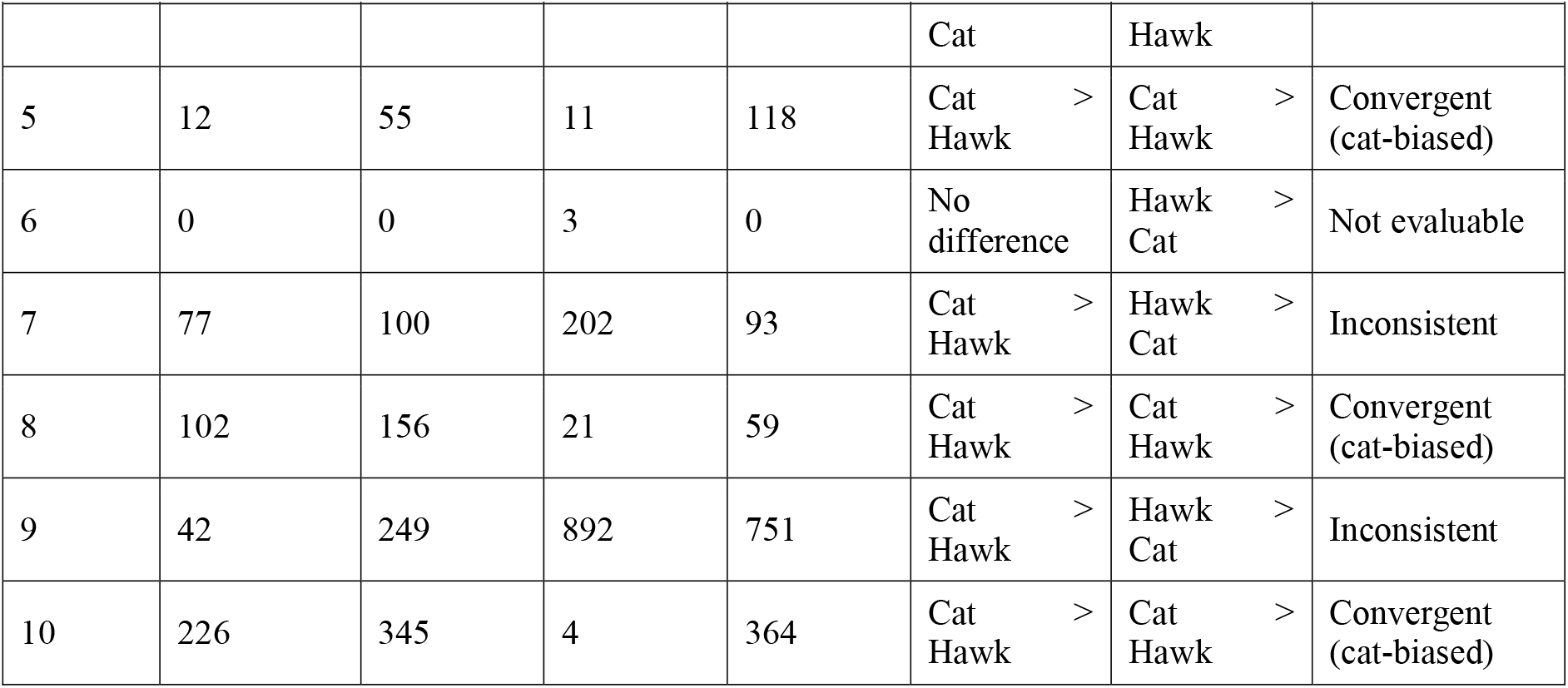
Total call frequency by predator type and individual, direction of the cat-versus hawk-exposure difference, and cross-individual convergence, for each of the 10 quantitatively characterized call types.

Three call types (5, 8, and 10) showed a consistent cat-biased pattern in both individuals. One call type (2) showed a consistent hawk-biased pattern in both individuals. The remaining six call types showed directions that differed between individuals.

### Shift in call-type composition before versus after predator exposure

The relative composition of the vocal repertoire differed markedly between the pre-exposure and post-exposure phases. For each of the four individual-by-predator combinations, a chi-square test of independence was performed on the contingency table of call type (10 levels) by phase (pre-versus post-exposure). All four tests were highly significant (Table 2).

**Table 2.** Chi-square tests comparing call-type composition between the pre-exposure and post-exposure phases, for each individual-by-predator combination.

| Combination | Chi-square | p-value | Pre-exposure n | Post-exposure n |
| --- | --- | --- | --- | --- |
| Ind. 0.0.0 – Hawk | 75 | < 0.0001 | 931 | 11 |
| Ind. 0.0.0 – Cat | 49.9 | < 0.0001 | 1007 | 78 |
| Ind. 795 – Hawk | 77.7 | < 0.0001 | 1114 | 483 |
| Ind. 795 – Cat | 112.3 | < 0.0001 | 1666 | 221 |

Inspection of the underlying proportions (Fig 2) reveals a pattern consistent across all four combinations: call type 9 increased its relative share of total calling after predator exposure ended, regardless of individual or predator class (Individual 0.0.0-Hawk: 4.3% to 18.2%; Individual 0.0.0-Cat: 22.5% to 28.2%; Individual 795-Hawk: 52.8% to 62.9%; Individual 795-Cat: 36.7% to 63.3%). This four-way convergence was the most consistent directional pattern observed in the present dataset. Several other call types (2, 3, 4, and 8) diminished or disappeared from the post-exposure repertoire in most combinations, suggesting an overall narrowing of the active vocal repertoire once the immediate predator stimulus had ended. In the combination with the fewest post-exposure calls (Individual 0.0.0-Hawk, n = 11), individual proportions are correspondingly unstable; the type 9 pattern is more robust in the two combinations with larger post-exposure totals (Individual 795-Hawk, n = 483; Individual 795-Cat, n = 221).

**Fig 1.**
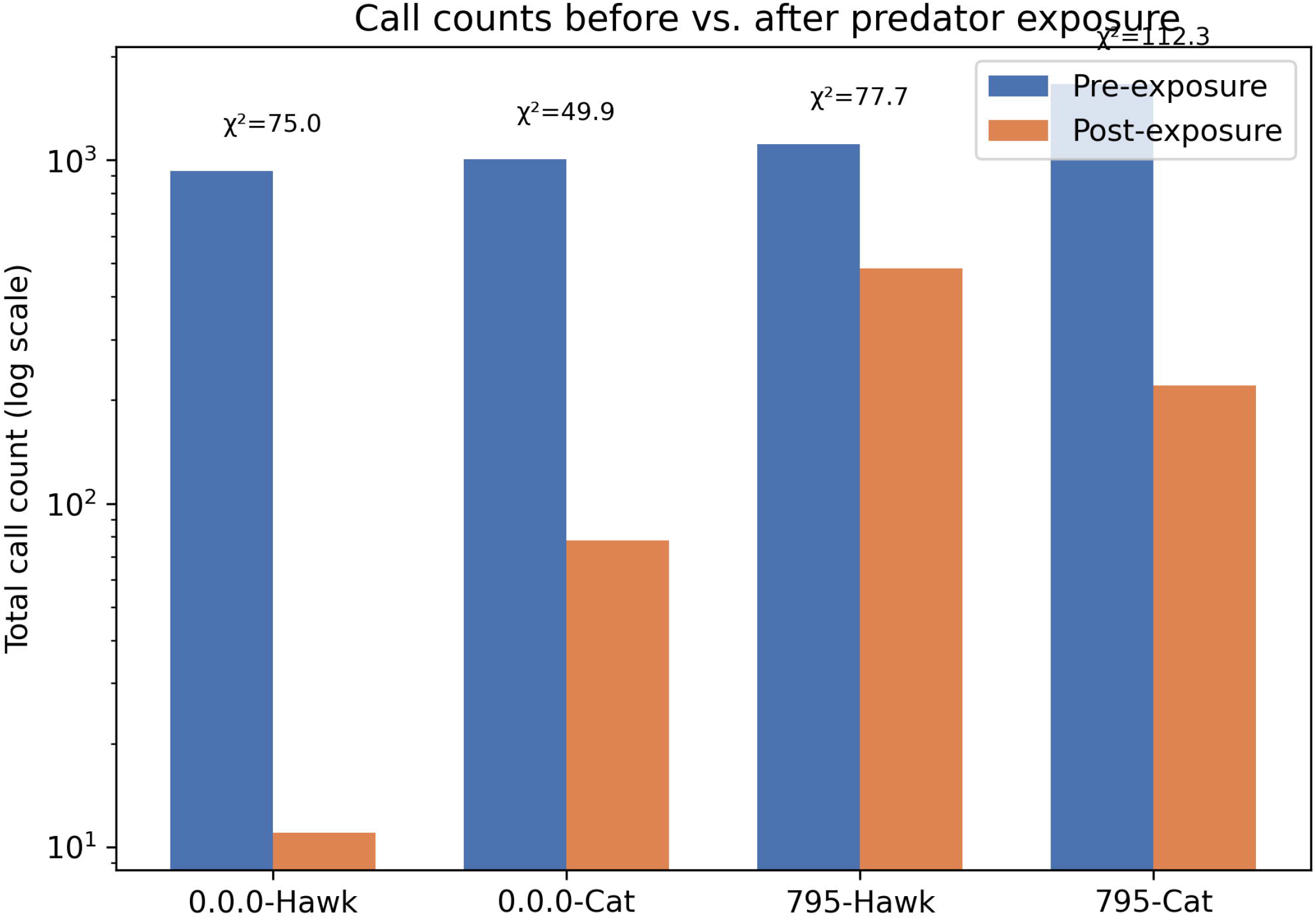
Total call counts before and after predator exposure for each individual-by-predator combination (log scale), with chi-square and p-values for the composition-shift test in Table 2.

**Fig 2.**
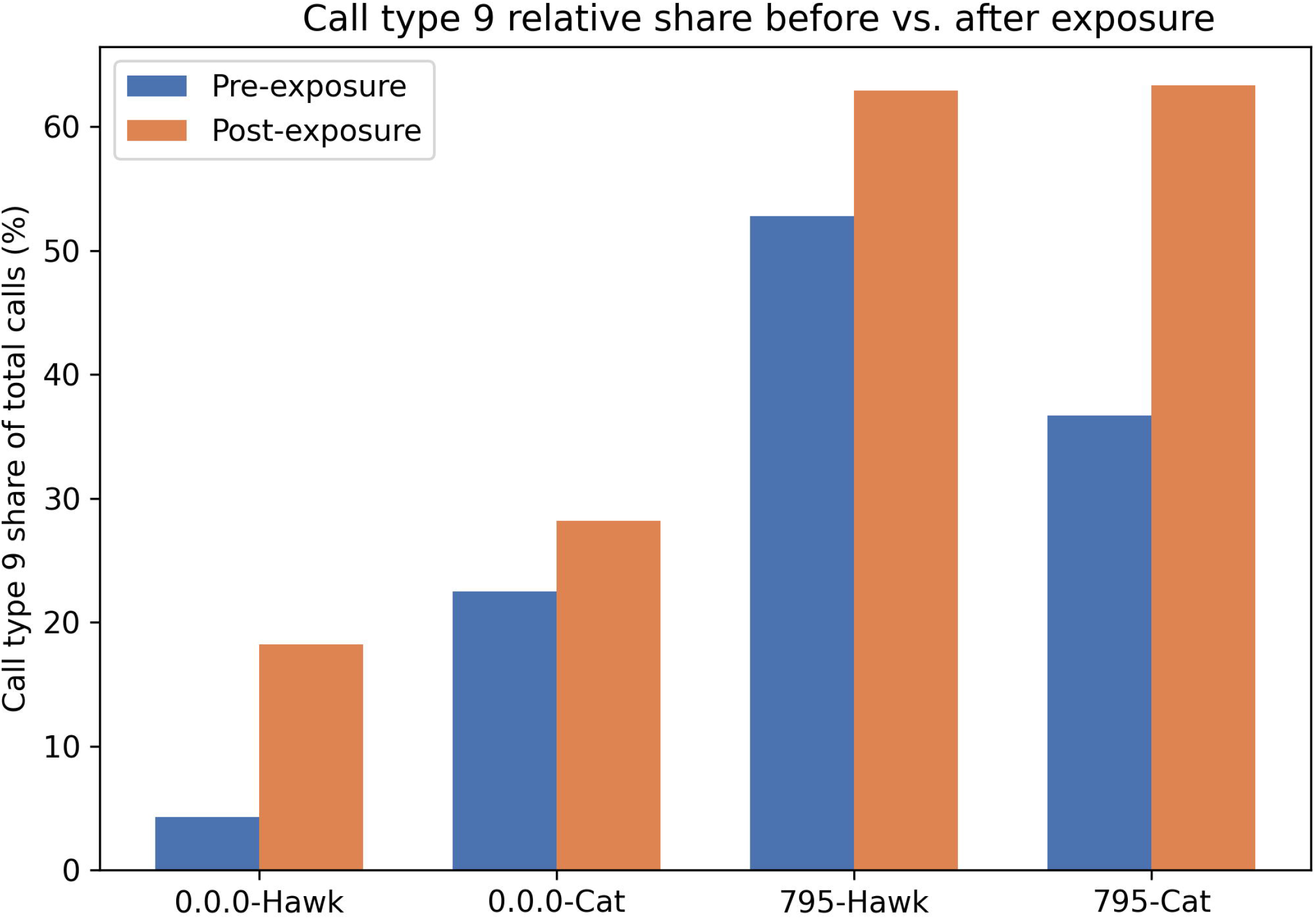
Call type 9’s share of total calls before (blue) and after (orange) predator exposure, shown separately for each individual-by-predator combination. This call type consistently increased in relative share after exposure across all four combinations.

### Predator-type composition differences and their acoustic correlates

A chi-square test of independence was performed on the call-type-by-predator contingency table separately for each individual, and standardized residuals were examined to identify which call types drove any overall difference (Table 3; Fig 3).

**Table 3.** Standardized residuals for call-type composition under hawk versus cat exposure, by individual. Positive residuals indicate overrepresentation under hawk exposure; negative residuals indicate overrepresentation under cat exposure. Values exceeding |1.96| are marked with **.

| Call type | Ind. 0.0.0 residual | Ind. 795 residual | Convergent | Direction |
| --- | --- | --- | --- | --- |
| 1 | 6.53 ** | -0.68 | No | -- |
| 2 | 3.11 ** | 3.59 ** | Yes | Hawk-biased |
| 3 | 3.26 ** | 1.22 | Directional only | Hawk-biased trend |
| 4 | 6.69 ** | -1.95 | No | -- |
| 5 | -3.43 ** | -6.26 ** | Yes | Cat-biased |
| 7 | -0.58 | 5.74 ** | No | -- |
| 8 | -1.63 | -2.59 ** | Directional only | Cat-biased trend |
| 9 | -8.02 ** | 5.05 ** | No | -- |
| 10 | -2.42 ** | -12.68 ** | Yes | Cat-biased |

**Fig 3.**
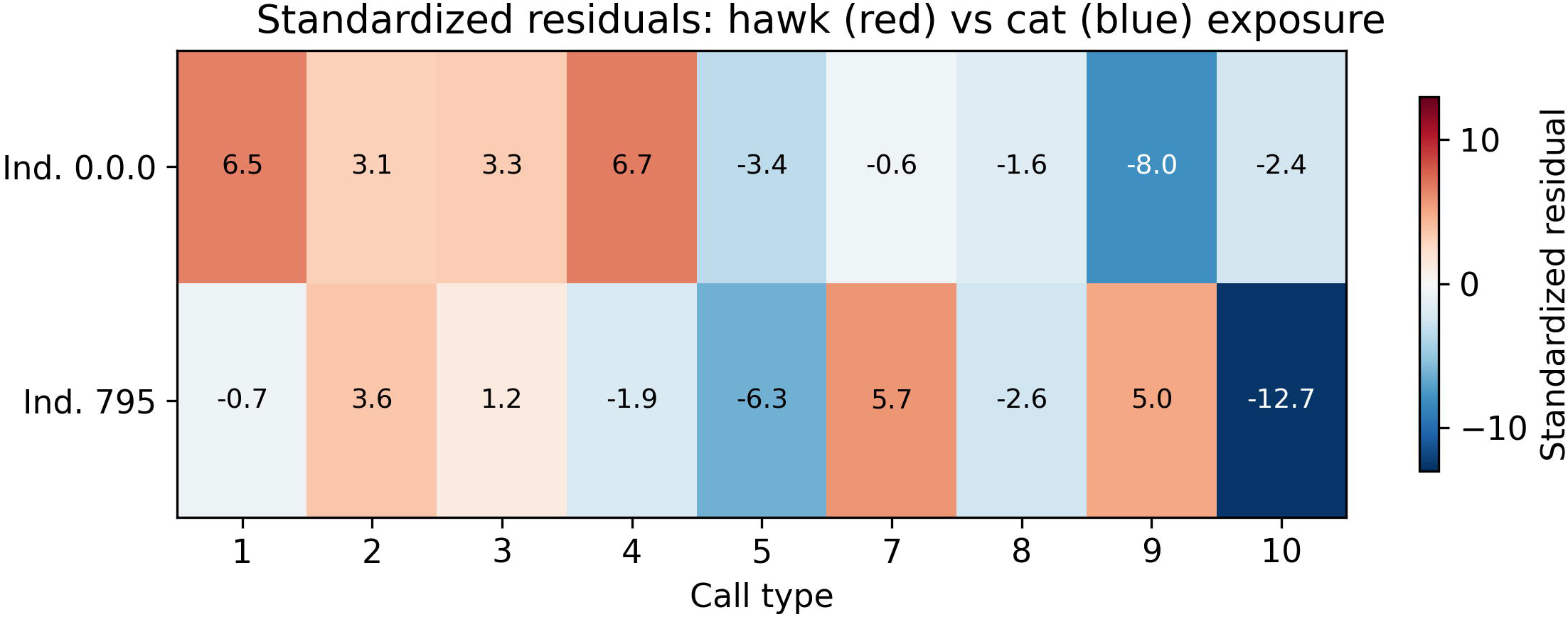
Heatmap visualization of Table 3: standardized residuals for call-type usage under hawk versus cat exposure, by individual. Red indicates hawk-biased overrepresentation; blue indicates cat-biased overrepresentation.

Composition differed significantly between hawk and cat exposure in both individuals (Individual 0.0.0: chi-square = 359.8, df = 8, p < 0.0001; Individual 795: chi-square = 527.7, df = 9, p < 0.0001). Call types 5 and 10 were significantly cat-biased in both individuals, and call type 2 was significantly hawk-biased in both individuals, corroborating the simpler total-count comparison above with a full compositional analysis.

As an exploratory follow-up, we examined whether the acoustic structure of a call type predicted its degree of hawk-versus-cat bias. For each of the ten quantified call types, a log2 ratio of pooled cat-exposure to hawk-exposure call counts was correlated with each acoustic parameter (Table 4; Fig 4).

**Table 4.** Correlation between call-type acoustic parameters and predator-type bias (log2 ratio of cat-to hawk-exposure call counts, pooled across both individuals; n = 10 call types).

| Acoustic parameter | Pearson r | p-value | Spearman rho | p-value |
| --- | --- | --- | --- | --- |
| Duration | -0.580 | 0.079 | -0.498 | 0.143 |
| Dominant frequency | -0.229 | 0.525 | -0.224 | 0.533 |
| Frequency range | -0.562 | 0.091 | -0.539 | 0.108 |
| Low frequency | 0.014 | 0.970 | -0.006 | 0.987 |
| High frequency | -0.505 | 0.136 | -0.673 | 0.033 |

**Fig 4.**
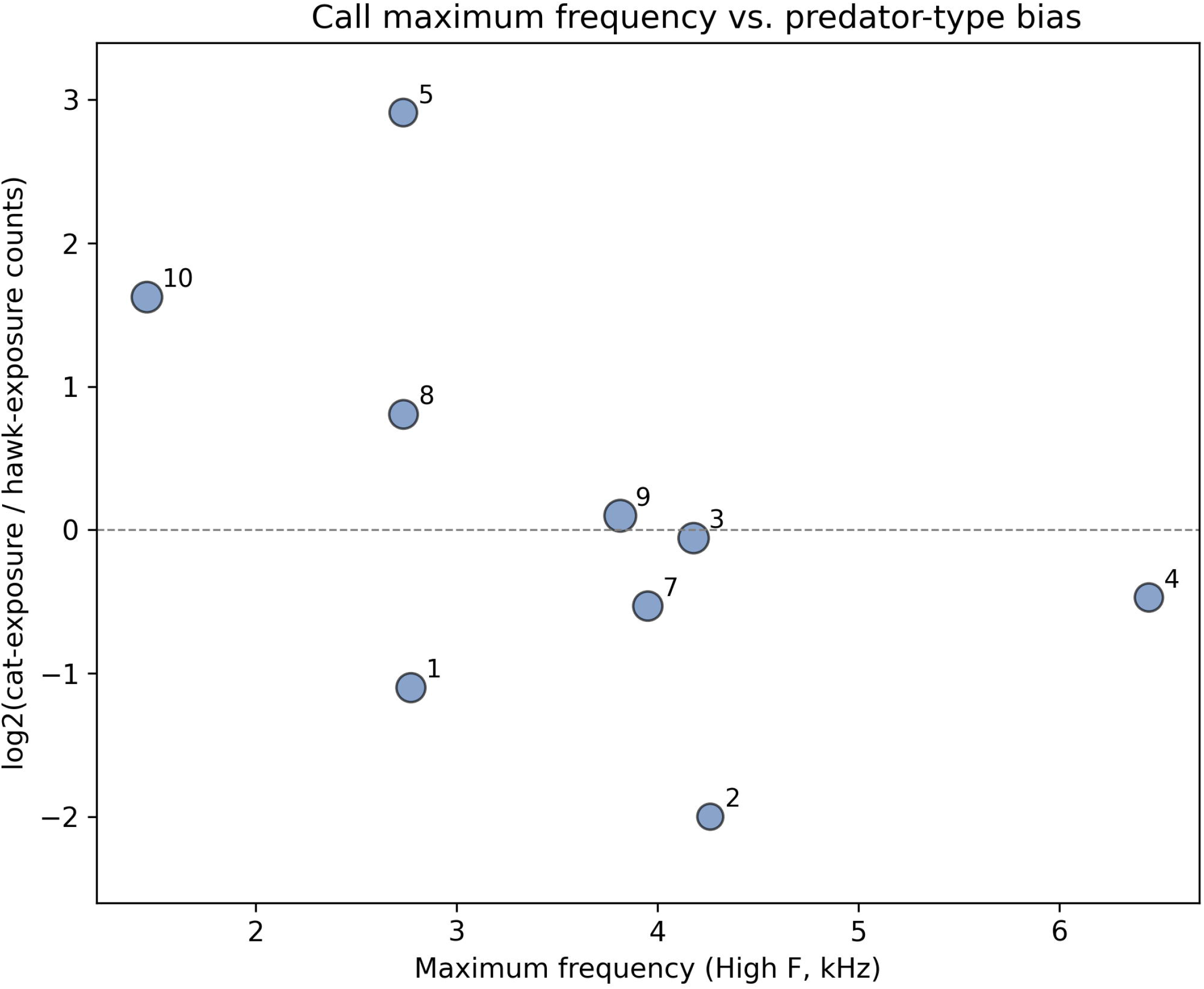
Relationship between a call type’s maximum frequency (High F) and its predator-type bias (log2 ratio of cat-to hawk-exposure counts, pooled across individuals). Marker size reflects the total number of calls recorded for that type (log scale).

Maximum frequency was significantly correlated with predator-type bias by Spearman’s rank correlation (rho = -0.673, p = 0.033), indicating that call types with a higher maximum frequency tended to be relatively hawk-biased. Given that five acoustic parameters were tested, this result did not survive a conservative Bonferroni correction (alpha = 0.01) and should be treated as hypothesis-generating rather than confirmatory.

## Discussion

This pilot case-series provides preliminary evidence that specific, spectrographically defined call types within the Japanese quail vocal repertoire are differentially associated with predator class, and that the composition of the repertoire narrows toward a small number of dominant types once a threat has passed. The convergent cat-biased calling observed for types 5, 8, and 10 in both recorded individuals, together with the convergent hawk-biased pattern for type 2, is consistent with the broader principle, established for domestic fowl [1,2] and functionally validated through playback [3], that avian vocal repertoires encode distinctions among external referents through structurally and behaviourally specific call types rather than through a single generic alarm response.

The consistent post-exposure increase in the relative share of call type 9, replicated across both individuals and both predator classes, represents the most robust convergent pattern in this dataset, more robust than any single call type’s absolute-frequency bias toward one predator class. Given its moderate duration and comparatively wide, mid-to-high frequency range, type 9 is a plausible candidate for a post-threat reassessment or alerting signal, broadly consistent with the general principle that avian repertoires narrow toward a small set of dominant, context-specific signals once acute threat has passed [1].

A plausible behavioural analogue for this candidate signal is offered by the vigilance trill described for European quail, which is reported to shift gradually from a short, high-pitched trill to a longer, lower-pitched trill as the bird transitions from active defence toward a watchful, vigilant state following an encounter [7]. The post-exposure increase in call type 9 observed here is broadly consistent with the existence of a distinct, transitional vigilance-related signal following the acute phase of a predator encounter in this genus more generally, although direct acoustic correspondence between call type 9 and the European-quail vigilance trill has not been established and would require formal comparison of the underlying sonograms. A companion analysis of the same underlying dataset (in preparation; details available from the corresponding author on request) undertakes exactly this comparison, testing the correspondence between each of the present study’s ten quantitatively characterized call types — including call type 9, there labelled JQ-C — and the named call catalogue described in [7], using both sonogram morphology and a quantitative acoustic-distance index.

These observations should be interpreted cautiously against the existing quail literature. The vocal repertoire of European quail (Coturnix coturnix coturnix) has been described in comparable detail, comprising 24 calls organized into four functional groups (social cohesion during routine activity, strong interattractive tendencies in reproductive contexts, intraspecific aggressive or defensive tendencies, and strong defensive tendencies in an interspecific context, the last of which encompasses several distinct alarm-, escape-, and vigilance-related calls), with an almost total overlap reported between the European and Japanese quail repertoires [7]. Within that four-group scheme, calls in the interspecific defensive category were not further differentiated by predator identity, consistent with the specific gap addressed by the present study. Two caveats apply to this comparison. First, Guyomarc’h and Guyomarc’h [7] directly characterized the European subspecies (C. c. coturnix); their conclusion of near-total overlap with Japanese quail (C. c. japonica) was based on comparative field and laboratory observation rather than an equivalently detailed spectrographic classification of the Japanese subspecies itself, so it should not be treated as establishing a validated one-to-one correspondence between subspecies. Second, and for the same reason, direct correspondence between our 18 call types and their 24 described European-quail calls cannot be established from textual description alone; a side-by-side spectrographic comparison, using the original sonogram images from both studies, would be required to test this correspondence rigorously and is identified here as a priority for future work; a companion analysis addressing this question (in preparation) is available from the corresponding author on request. Separately, acoustic parameters describing call timing have been shown to discriminate reliably between individuals and between subspecies [8].

The acoustic-correlate finding only partially aligns with the classic prediction that aerial-predator alarm calls should be narrow-band, high-frequency signals that are difficult for the predator to localize (the ‘seet call’ hypothesis [15]): while our hawk-biased call types did tend toward higher maximum frequency, they also trended toward wider, not narrower, frequency range and longer duration, which is not fully consistent with a pure localization-difficulty account. This discrepancy may reflect genuine species-specific differences, the small number of call types and individuals examined, or the fact that call types in this dataset are not yet confirmed to function specifically as anti-predator alarm signals rather than general arousal-linked vocalizations.

Complementary exploratory analyses of social-context manipulations (mirror exposure and a concealed, audible-only conspecific) are reported in the supporting information. Briefly, call-type composition under mirror exposure resembled that under live-conspecific exposure more closely than that under solitary conditions in three of four combinations, and composition under the concealed-conspecific condition remained more similar to the mirror-alone pattern than to the fully visible conspecific pattern in all four combinations. A directly analogous design in the domestic chicken found that roosters emitted alarm calls toward a real conspecific but not toward their own mirror reflection, even when a real conspecific was concealed behind the mirror, while nonetheless failing a classic mark test [11]; our finding that the concealed-conspecific condition remained acoustically anchored to the mirror-alone pattern is broadly consistent with this precedent. These patterns should not be interpreted as evidence for or against mirror self-recognition, which requires a dedicated mark-test paradigm [16]; they are reported as tentative, case-based observations that may inform future study designs (S1 File).

Evaluated against the STRANGE framework [14], the present sample is limited in size (n = 2) and, pending full documentation of housing and rearing history, its representativeness relative to the broader Japanese quail population cannot be established. The findings reported here should accordingly be regarded as case-based hypotheses rather than generalizable population-level effects.

## Limitations

This study has six limitations that must be addressed before its findings can be generalized. First, the sample consists of only two individuals; however, each individual contributed several thousand classified vocalizations across repeated predator-exposure trials, and the same directional effects (call-type-specific predator bias, post-exposure reorganization) replicated independently in both individuals and both predator classes — a repeated-measures, single-case-series design comparable to small-N designs used elsewhere in behavioural science to establish reproducible within-subject effects before a fully powered population-level study is undertaken. We nonetheless evaluate this sample, under the STRANGE framework, as a source of unresolved potential sampling bias, and the convergence patterns reported should be read as case-based, reproducible observations rather than statistically generalizable population-level effects; a confirmatory follow-up study with a minimum of 8–10 individuals per predator condition, informed by the effect sizes observed here, is planned to formally test generalizability at the population level. Second, unobserved cells in the underlying dataset were coded as missing rather than as confirmed zero counts, introducing ambiguity that should be resolved against the original experimental protocol. Third, the present results characterize call production only; whether these call types carry functionally referential information remains to be tested with playback experiments [3]. Fourth, total call counts were substantially higher in the pre-exposure phase than in the post-exposure phase for both individuals, opposite in direction to the within-exposure suppression and post-exposure rebound previously reported for pooled separation calls in this species and laboratory [12]; it must be confirmed whether the pre- and post-exposure observation windows were of equal duration. Fifth, the acoustic-correlate analysis tested five parameters without correction for multiple comparisons; only maximum frequency reached nominal significance, and this result should be treated as exploratory. Sixth, although the 18-call-type classification was confirmed by blind review from a second, independent observer, we did not compute a formal inter-rater reliability statistic (e.g., Cohen’s kappa), and readers should treat the size of the described repertoire as a qualitatively cross-validated, exploratory estimate rather than a statistically quantified or definitively established taxonomy; independent, quantitative support for a subset of these call types against an established congeneric reference catalogue is provided in a companion sonogram-comparison analysis (in preparation; available from the corresponding author on request).

## Conclusions

Under a pilot case-series design, two Japanese quail showed reproducible, predator-type-associated shifts in the use of specific call types within a broader 18-type vocal repertoire, together with a consistent narrowing and reorganization of the repertoire following predator exposure. These findings justify a fully powered follow-up study, incorporating a larger number of individuals, complete STRANGE-framework reporting, an explicit accounting of missing observation periods, and playback-based validation of functional referentiality, to determine whether the Japanese quail vocal repertoire encodes predator-class-specific information in a manner analogous to that established in the domestic chicken.

## Supporting information

cover letter

## Acknowledgments

We thank Sonja Hillemacher for her valuable advice and materials during the design of the predator-exposure and mirror-audience experiments, and Ju Hyun Lee for providing the software and methodology used for vocalization analysis.

## Supporting information

S1 File. Extended analyses of social-context (mirror and concealed-conspecific) effects on call-type composition. Contains the full contingency tables and chi-square results for social contexts B, C, and D referenced in the Materials and Methods and Discussion.

