## Supplementary material for "Predator type shapes call-type composition and post-threat vocal dynamics in Japanese quail (Coturnix japonica): a pilot case-series": cover letter

Chunghyun Kim, DVM, PhD

Department of Animal Health and Welfare

College of Biohealth Sciences, Hoseo University

Asan, Republic of Korea

Email: [corresponding author email]

2026.9.6.

Dear Editors,

Please consider our manuscript, “Predator type shapes call-type composition and post-threat vocal dynamics in Japanese quail (Coturnix japonica): a pilot case-series,” for publication in PLOS ONE as a Research Article.

Japanese quail possess an extensive vocal repertoire, but whether individual call types are differentially recruited according to predator identity, and how that repertoire reorganizes once a threat has passed, has not previously been examined systematically in this species. Using a repeated-measures pilot case-series design (two individuals, each contributing several thousand classified vocalizations across repeated hawk and cat exposure trials), we show that call-type composition differs significantly by predator class in both individuals, with convergent cat- and hawk-biased call types, and that one call type consistently increases its post-exposure share across all four individual-by-predator combinations — a pattern we relate to vigilance-related signalling described for the congeneric European quail. We report these as case-based, hypothesis-generating observations given the sample size, and detail a concrete follow-up design (minimum 8–10 individuals per condition) that these pilot results are intended to justify.

This work relates most directly to foundational studies of functionally referential and audience-modulated alarm calling in the domestic chicken, and to the descriptive vocal repertoire established for the congeneric European quail; full details are cited in the Introduction and Discussion.

This is a Research Article. We have had no prior interactions with PLOS regarding this manuscript.

Related manuscript disclosure: a companion manuscript from our group, analyzing the same underlying acoustic dataset from a different angle (a sonogram-based cross-species comparison of the resulting call types against an existing congeneric vocal catalogue, rather than the predator-composition analysis reported here), is currently in preparation and, separately, under consideration at another journal. The two manuscripts address distinct questions, use independent call-type labelling schemes, and are each designed to stand as an independent unit; we are glad to provide a copy of the companion manuscript to the editor on request.

Suggested Academic Editors: we suggest an editor with expertise in avian bioacoustics, galliform/poultry behavior, or predator-elicited alarm signalling; we did not identify a specific conflict-free individual from the current PLOS ONE editorial board with sufficient confidence to name one, and defer to the editorial office's assignment.

We have no opposed reviewers to list.

All authors have approved this submission. We have no conflicts of interest to declare. Thank you for considering our manuscript.

Sincerely,

Chunghyun Kim and Goun Park (co-corresponding authors)

on behalf of all co-authors (Kayeon Ham, Seeun Park, Goun Park, Chunghyun Kim)
